# Direct Pathway Neurons in Mouse Dorsolateral Striatum *In Vivo* Receive Stronger Synaptic Input than Indirect Pathway Neurons

**DOI:** 10.1101/651711

**Authors:** Marko Filipović, Maya Ketzef, Ramon Reig, Ad Aertsen, Gilad Silberberg, Arvind Kumar

## Abstract

Striatal projection neurons, the medium spiny neurons (MSNs), play a crucial role in various motor and cognitive functions. MSNs express either D1 or D2 type dopamine receptors and initiate the direct-pathway (dMSNs) or indirect pathways (iMSNs) of the basal ganglia, respectively. dMSNs have been shown to receive more inhibition than iMSNs from intrastriatal sources. Based on these findings, computational modelling of the striatal network has predicted that under healthy conditions dMSNs should receive more excitatory input than iMSNs. To test this prediction, we analyzed *in vivo* whole-cell recordings from dMSNs and iMSNs in healthy and dopamine-depleted (6OHDA) anaesthetized mice. By comparing their membrane potential fluctuations, we found that dMSNs exhibited considerably larger membrane potential fluctuations over a wide frequency range. Furthermore, by comparing the spike-triggered average membrane potentials, we found that dMSNs depolarized towards the spike threshold significantly faster than iMSNs did. Together, these finding corroborate the theoretical prediction that direct-pathway MSNs receive stronger input than indirect-pathway neurons. Finally, we found that dopamine-depleted mice exhibited no difference between the membrane potential fluctuations of dMSNs and iMSNs. These data provide new insights into the question how a lack of dopamine may lead to behavior deficits associated with Parkinson’s disease.

**Significance statement:** The direct and indirect pathways of the basal ganglia originate from the D1 and D2 type dopamine receptor expressing medium spiny neurons (dMSNs and iMSNs), respectively. To understand the role of the striatum in brain function and dysfunction it is important to characterize the differences in synaptic inputs to the two MSN types. Theoretical results predicted that dMSNs should receive stronger excitatory input than iMSNs. Here, we studied membrane potential fluctuation statistics of MSNs recorded *in vivo* in anaesthetized mice and found that dMSNs, indeed, received stronger synaptic input than iMSNs. We corroborated this finding by spike-triggered membrane potential analysis, showing that dMSNs spiking required more synaptic input than iMSNs spiking did, as had been predicted by computational models.

## Introduction

The striatum is the largest nucleus in the basal ganglia (BG) and acts as its main input structure. GABAergic medium spiny neurons (MSNs) are the striatal projection neurons and constitute about 95% of the striatal neuronal population. D1 type dopamine receptor expressing MSNs (dMSNs) project to the substantia nigra pars reticulata and globus pallidus interna and constitute the ‘direct pathway’, whereas D2 type dopamine receptor expressing MSNs (iMSNs) project to the globus pallidus externa and constitute the ‘indirect pathway’. A balance in the activity of the two pathways is essential for correct functioning of the BG, and is disrupted in BG-related pathologies such as Parkinson’s disease (PD). To understand how the direct and indirect pathways shape BG function, we need to quantify both the upstream excitatory inputs into the striatum and the recurrent inhibitory connections within and between dMSNs and iMSNs.

The dMSNs and iMSNs differ in their connectivity: iMSN to dMSN connectivity (13%) is much higher than dMSN to iMSN (4.5%), whereas dMSN to dMSN connectivity (7%) is much lower than iMSN to iMSN (23%) (Taverna et al., 2008; Planert et al., 2010). Moreover, GABAergic fast-spiking interneurons (FSIs) connect preferentially to dMSNs compared to iMSNs (53% vs. 36%) (Gittis et al., 2010). That is, dMSNs receive overall more inhibition than iMSNs. Despite these differences, both dMSNs and iMSNs exhibit similar average activity in awake behaving animals (Cui et al., 2013; Sippy et al., 2015).

Using a computational model we recently predicted that dMSNs should receive stronger excitatory input than iMSNs (either through more synapses, stronger synapses, or stronger input rates and/or correlations), so that both dMSNs and iMSNs may have comparable firing rates (Bahuguna et al., 2015). Recent *ex vivo* recordings suggest that cortico-striatal synapses on dMSNs may be stronger than those on iMSNs (Parker et al., 2016) (however, see Lei et al. (2004); Kress et al. (2013); Doig et al. (2010); Deng et al. (2015)). While this data supports the theoretical predictions, it is well known that *in vivo* synaptic conductances can be very different from *ex vivo* measurements (Destexhe et al., 2003).

Even though it is hard to estimate the full strength and numbers of individual excitatory synapses impinging on dMSNs and iMSNs experimentally, a relative difference in the total input to the two neuron types can be estimated by analyzing *in vivo* intracellular membrane potential fluctuations. In particular, the variance (or the spectral power) of the membrane potential fluctuations is proportional to the square of the synaptic strength (Kuhn et al., 2004). That is, by comparing the spectra of sub-threshold membrane potential *in vivo* we can test whether dMSNs indeed receive stronger excitatory input than iMSNs, as was theoretically predicted (Bahuguna et al., 2015).

Therefore, we recorded and analyzed the *in vivo* membrane potentials of dMSNs and iMSNs from healthy and dopamine-depleted anaesthetized mice using whole-cell patch clamp recordings. These neurons exhibited alternating periods of high and low activity (called up- and down-states, respectively), characteristic of recordings in animals under ketamine-induced anaesthesia (Wilson and Kawaguchi, 1996). We found that dMSNs exhibited higher spectral power in their up-states than iMSNs over a wide range of frequencies in healthy mice. In addition, bilateral whisker stimulation in healthy animals showed that sensory inputs evoked larger responses in dMSNs than in iMSNs. Despite these differences, the membrane time constants of the two MSN types were not significantly different. According to linear systems theory, stronger membrane potential fluctuations are indicative of stronger synaptic inputs and/or higher input correlations (Kuhn et al., 2004). Finally, we found that dopamine depletion abolished the difference in spectral power of up-state membrane potential fluctuations between dMSNs and iMSNs, highlighting the role of dopamine in maintaining the activity balance between the direct and indirect pathways.

Thus, our study provides the first experimental *in vivo* evidence of stronger synaptic input to the direct-pathway of the mouse dorsolateral striatum, and demonstrates that this difference is attenuated in dopamine-depleted animals.

## Methods

### Experimental Methods

#### Ethics approval

All experiments were performed according to the guidelines of the Stockholm municipal committee for animal experiments under an ethical permit to G.S. (N12/15). D1-Cre (EY262 line) or D2-Cre (ER44 line, GENSAT) mouse line were crossed with the Channelrhodopsin (ChR2)-YFP reporter mouse line (Ai32, Jackson laboratory) to induce expression of ChR2 in either dMSNs or iMSNs, respectively. Mice of both sexes were housed under a 12-hour light-dark cycle with food and water ad libitum. All experiments were carried out during the light phase.

#### 6OHDA lesioning

Mice (12 males and females 8-10 weeks of age) were anesthetized with isoflurane and mounted in a stereotaxic frame (David Kopf Instruments, Tujunga, California). The mice received one unilateral injection of 1 µL of 6OHDA-HCl (3.75 µg/µL dissolved in 0.02 % ascorbic acid) into the medial forebrain bundle (MFB), according to the following coordinates (Paxinos and Franklin, 2004): antero-posterior −1.2 mm, medio-lateral 1.2 mm and dorso-ventral −4.8 mm. After surgery, all mice were injected with Temgesic (0.1 mg/kg, Reckitt Benckiser, Berkshire, England) and allowed to recover for at least 2 weeks. Sham and unlesioned mice (n = 21 of both sexes) served as controls, their data were pooled after no differences were found between the groups. Only 6OHDA injected mice that showed rotational behavior (Santini et al., 2007) were used in our experiments.

#### In vivo recordings

Experiments were conducted as described previously (Reig and Silberberg, 2014; Ketzef et al., 2017). Briefly, 2-3 weeks post-lesioning, mice were anesthetized by intraperitoneal (IP) injection of ketamine (75 mg/kg) and medetomidine (1 mg/kg) diluted in 0.9 % NaCl. To maintain mice under anesthesia, a third of the dose of ketamine was injected intraperitonally approximately every 2 hours or in case the mouse showed response to pinching or changes in EcoG patterns. Mice were tracheotomized, placed in a stereotactic frame, and received oxygen enriched air throughout the recording session. Core temperature was monitored with a feedback-controlled heating pad (FHC) and kept on 36.5±0.5 °C. Patch clamp recordings were performed in the dorsolateral striatum since the sensory and motor areas project topographically onto it (McGeorge and Faull, 1989). The skull was exposed and a craniotomy was drilled (Osada success 40) 3.5-4 mm lateral to the bregma, and the dura was removed. Patch pipettes were pulled with a Flaming/Brown micropipette puller P-1000 (Sutter Instruments). Pipettes (7-10 MΩ, borosilicate, Hilgenberg), back-filled with intracellular solution, were inserted with a ∼1500 mbar positive pressure to a depth of about 2 mm from the surface, after which the pressure was reduced to 30-35 mbar. The pipette was advanced in 1 µm steps in depth (35 degrees angle), in voltage clamp mode. When a cell was encountered, the pressure was removed to form a Gigaseal, followed by application of a ramp of increasing negative pressure until a cell opening was evident. Recording was performed in current clamp mode. Intracellular solution contained: 130 K-gluconate, 5 KCl, 10 HEPES, 4 Mg-ATP, 0.3 GTP, 10 Na2-phosphocreatine, and 0.2-0.3 % biocytin (pH = 7.25, osmolarity 285 mOsm). The exposed brain was continuously covered by 0.9 % NaCl to prevent drying. Signals were amplified using a MultiClamp 700B amplifier (Molecular Devices) and digitized at 20 kHz with a CED acquisition board and Spike 2 software (Cambridge Electronic Design).

#### Optogenetic identification of in vivo recorded neurons

To obtain “on line” identification of whole-cell recorded neurons, we used the optopatcher (Katz et al., 2013) (A-M systems, WA USA). Computer controlled pulses of blue light (7 mW LED, 470 nm, Mightex systems) were delivered through an optic fiber inserted into the patch-pipette while recording the responses in whole-cell configuration (Fig. 1*A*). Light steps (500 ms) were delivered every 2-5 seconds with increasing intensity between 20 to 100 % of full LED power (2.1 mW at the tip of the fiber). Positive cells responded to light stimulation by step-like depolarization with or without firing, whereas negative cells did not show any response (Fig. 1*B*, and see Ketzef et al. (2017) for full characterization).

**Fig 1.**
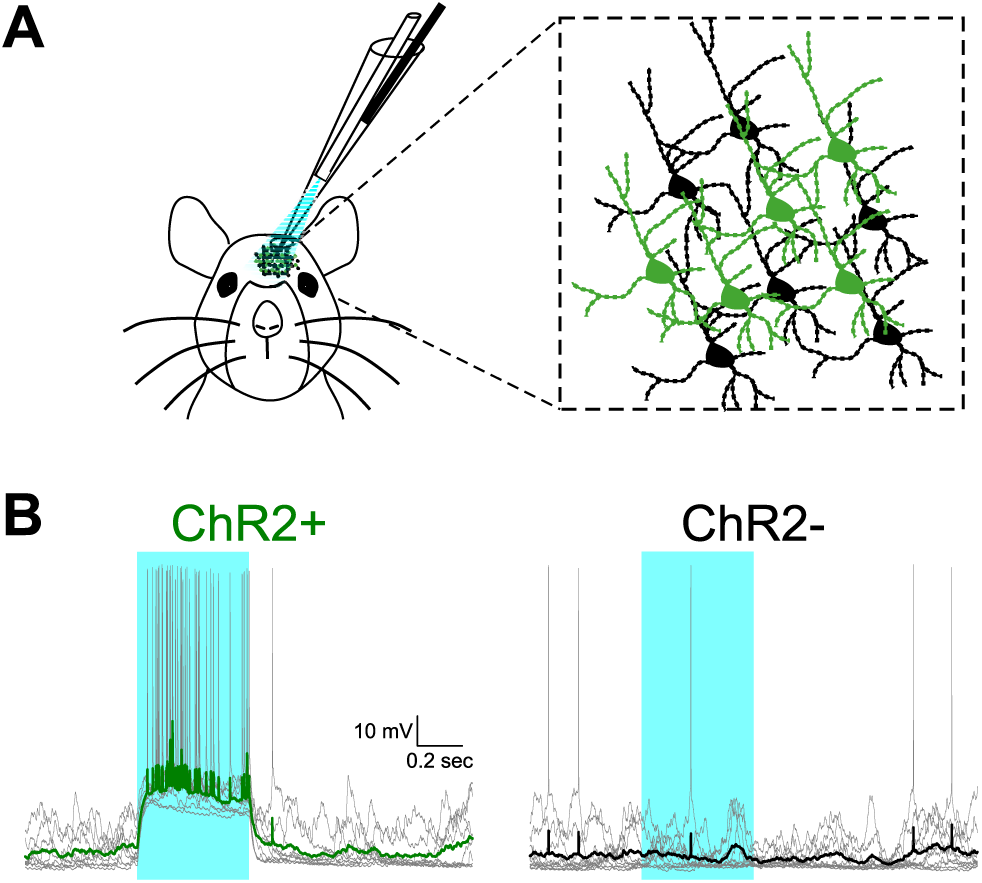
MSNs classification using the optopatcher. To facilitate the classification of MSNs as belonging to either the direct (dMSN) or indirect (iMSN) pathway in vivo, we utilized an optogenetic approach. In either D1-Cre or D2-Cre animals crossed with ChR2 reporter mouse, we selectively expressed ChR2 in dMSNs or iMSNs, respectively. Using the optopatcher, we could deliver focal light stimulation to the recorded cell and classify its identity ‘online’ during whole cell patch recordings. ***A*** Illustration of the experimental approach (*left*). In anesthetized mice, the optopatcher is introduced through the craniotomy. The optic fiber is inserted into the patch pipette and light application is focal. MSNs of both pathways are intermingled (*right*), positive cells (green) express ChR2 and YFP, whereas negative cells (black) do not. ***B*** Whole cell patch recording from positive (*left*) and negative (*right*) cells in a D2-ChR2 mouse. When the blue light is activated (470 nm, 0.5 s), positive cells depolarize immediately, whereas negative cells are not affected. Each example shows 10 repetitions (gray), overlaid by the average trace (green for positive and black for negative cells).

#### Whisker stimulation

Air puffs were delivered by a picospritzer (Picospritzer III, Parker Hannifin) through plastic tubes (1 mm diameter) positioned up to a centimeter from the mouse’s whiskers. Air puff stimulations (15 ms) were delivered at 0.2 Hz and at least 30 responses were acquired for each stimulation condition. The air pressure was set to 103.4-137.9 kPa (15-20 PSI).

### Data Analysis

#### Up- and down-state detection

For each membrane potential recording, we used a short time window (20-100 ms, depending on the noise level in the recording) to identify sudden transitions in the membrane potential with an amplitude sufficiently large to cross the cell-specific up-state or down-state thresholds. Upon detection of such a transition, we classified the following voltage period as an up-state or a down-state (Fig. 2*A*). The next sufficiently large membrane potential transition in the opposite direction marked the ending of that state. State thresholds were determined by finding the two main peaks of the bimodal voltage histogram of the entire trace, and by empirically adjusting these thresholds for the best detection rate (see also Léger et al. (2005) and Fucke et al. (2011)). In cells where the overall baseline voltage level fluctuated over time, we either used only the most stable section of the recording or discarded the entire recording alltogether. All states with a duration shorter than 40 ms were discarded from the analysis (Mukovski et al., 2007).

**Fig 2.**
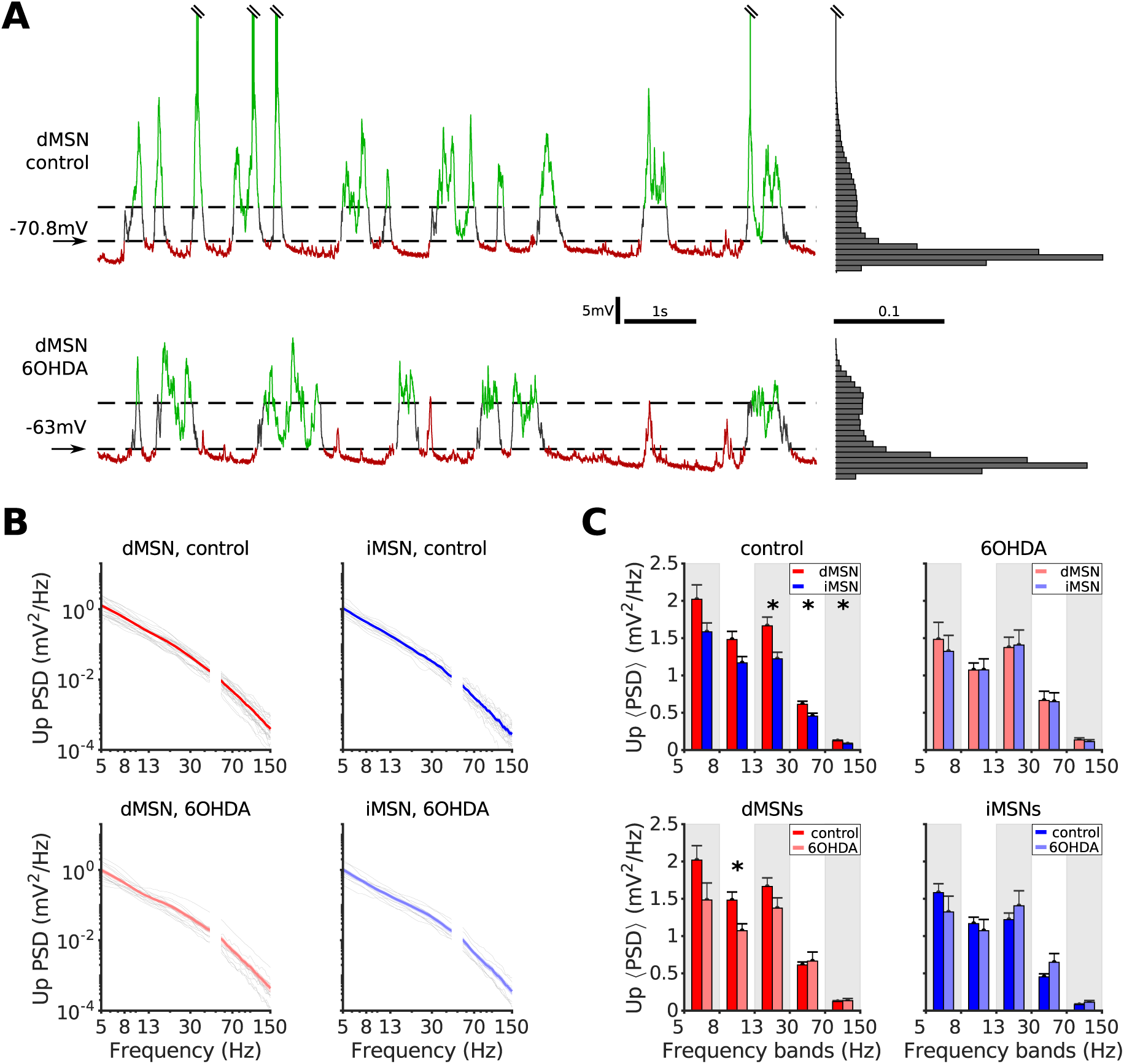
dMSNs carry more power than iMSNs during up-states in control conditions. ***A*** *Left*: Ten seconds of membrane potential recordings for a dMSN in control (upper trace) and 6OHDA (lower trace) conditions, exhibiting up- and down-states (green and red, respectively). Dashed lines represent the two cell-specific voltage thresholds used for state classification (see Methods). *Right*: Distributions of membrane potential values for the entire recordings of the two neurons shown at *left*. Note the characteristic bimodality of the up- and down-states. ***B*** Grand-average PSD estimates of up-states for all dMSNs and iMSNs in control (*top*, red and blue, respectively) and 6OHDA (*bottom*, light red and light blue, respectively) conditions. Grey traces represent average up-state PSD estimates of individual neurons. Frequencies between 45 and 55 Hz were removed to avoid power line contamination (see Methods). ***C*** Comparison of grand-average PSD estimates in different frequency bands. dMSNs exhibited higher power spectral density than iMSNs in control conditions in beta (*p* = 0.0135, *α*_*HB*_ = 0.0167), low-gamma (*p* = 0.0095, *α*_*HB*_ = 0.01), and high-gamma bands (*p* = 0.0103, *α*_*HB*_ = 0.0125; dMSN *n* = 26, iMSN *n* = 18 for all three bands), indicating either stronger or more frequent synaptic input. dMSNs also showed increased PSD in control versus 6OHDA for the 8-13 Hz band (*p* = 0.0087, *α*_*HB*_ = 0.01). Test statistics were corrected using the Holm-Bonferroni procedure.

For the purpose of characterizing the sub-threshold membrane potential dynamics analysis we excluded all up-states during which spiking occurred. Moreover, we also excluded a state from further analysis if one or more of the following criteria was met: (1) an up-state was interrupted by a down-state shorter than 25 ms (both states were discarded), (2) the mean membrane potential of a down-state exceeded the global average down-state potential for that cell by more than 3 %, (3) a recording artefact was present, or (4) a whisker stimulation trigger occurred either during the state or 200 ms preceding the state.

Finally, for all remaining states, we removed 5 % of the data, from the start of the state and before the end of the state, to minimize the impact of state transitions on the measured variables.

#### Power spectral density (PSD) estimation

The PSD estimate of an up-state membrane voltage trace was determined by first subtracting the mean potential from the remainder of the trace, and then by applying MATLAB’s periodogram function with Bartlett-Hann windowing. The minimal detectable frequency in individual up-state PSDs was set as the inverse of the duration of that state. For each cell, all such up-state PSD estimates were averaged to obtain a single power spectral density curve (Fig. 2*C*, gray traces). When comparing PSDs across cell groups (dMSNs vs. iMSNs), we constructed a grand-average PSD for each group by averaging over PSDs of individual cells (Fig. 2*C*, color traces). Frequencies below 5 Hz were disregarded because we observed only few up-states longer than 200 ms. Additionally, all frequency content between 45 and 55 Hz was removed to avoid power line contamination. We restricted the higher frequency range to 150 Hz, adopting this as the upper limit of the high-gamma band in our study.

We divided PSD estimates obtained in this manner into five frequency bands: sub-*α* (5-8 Hz), *α* (8-13 Hz), *β* (13-30 Hz), low-*γ* (30-70 Hz), and high-*γ* (70-150 Hz). To calculate the total power within any one frequency band for each cell, we isolated the section of interest of the PSD estimate and integrated the area under the curve.

Due to very low levels of spectral power during the down-states, the line noise power precluded any meaningful PSD comparison across the two cell groups.

#### Effective membrane time constant estimation

We estimated the effective membrane time constant *τ*_*m*_ from the *in vivo* membrane potential fluctuations by the following method. For a narrow enough voltage range, the membrane of a neuron can be approximated as a linear low-pass filter. Then, for that narrow voltage range, *τ*_*m*_ is directly proportional to the membrane capacitance and inversely related to the total membrane conductance. Thus, to minimize non-linear voltage-dependent effects, to account for the voltage dependence of *τ*_*m*_ (Kuhn et al., 2004), and to be able to treat the neuron membrane as a linear low-pass filter, we first binned the average membrane potentials of all states in 0.5 mV wide bins. Then we estimated the power spectral density of individual states belonging to each bin and averaged over the estimates in order to reduce noise, as explained in the previous section. Further noise reduction was achieved by smoothing the averaged PSD estimate with a Gaussian kernel, and the resulting curve was used to extract the cutoff frequency *f*_*c*_, calculated as the point where the maximal value of the smoothed PSD estimate fell to one half (−3 dB point, Fig. 3*A*). The initial effective membrane time constant 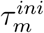 was then calculated as 1/(2*πf*_*c*_). We repeated this procedure for a series of narrow voltage ranges across different instances of up- and down-states within a single cell, in order to avoid non-linearities induced by large excursions of the effective membrane conductance.

**Fig 3.**
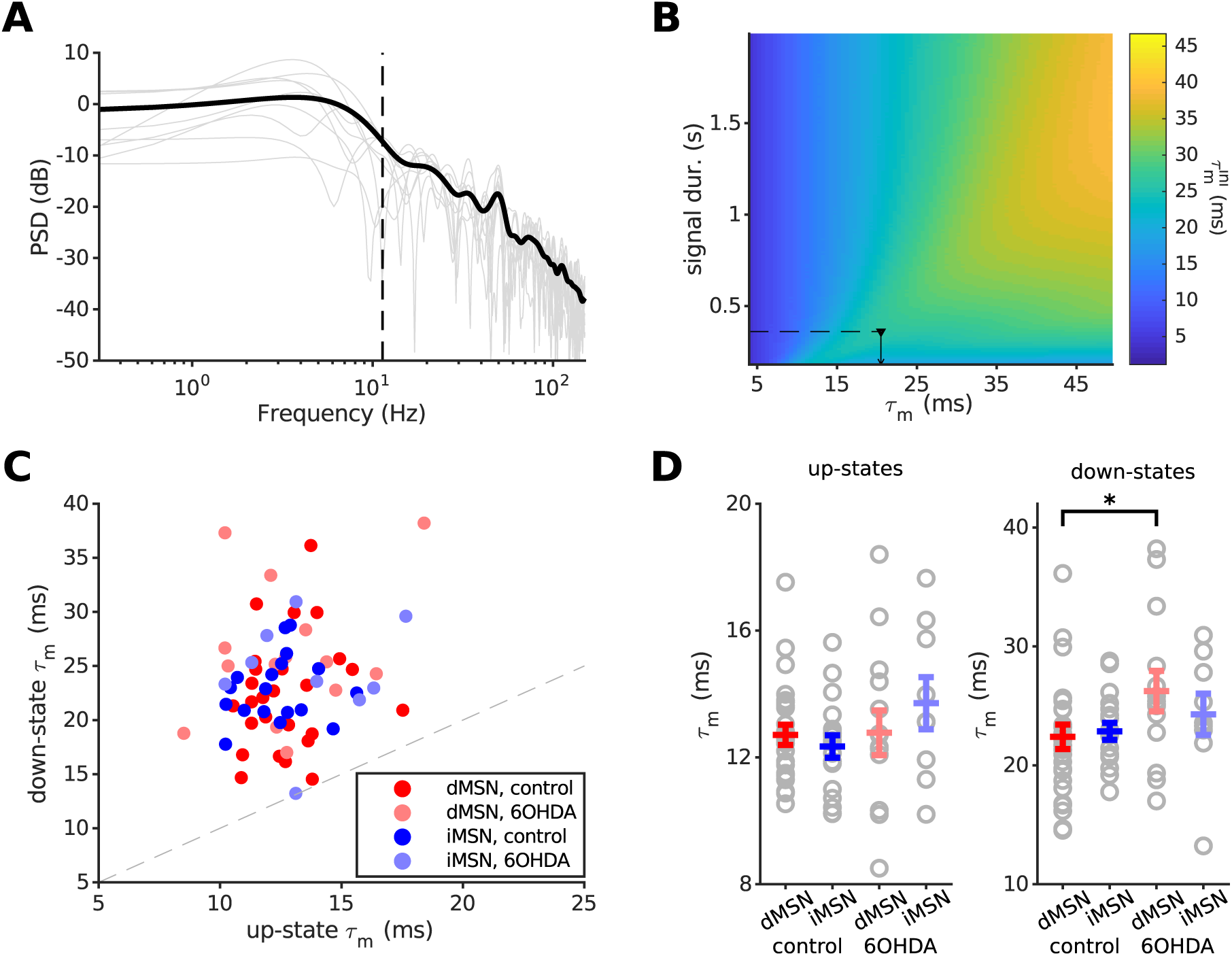
No difference in effective membrane time constant between dMSNs and iMSNs in up-states. ***A*** Example of the effective membrane time constant estimation in a dMSN, for all up-states with the mean membrane potential falling into a single 0.5 mV voltage bin. PSD estimates of individual up-states (grey) were averaged and smoothed (black trace), and the initial membrane time constant 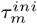 was estimated as the point where the maximal power decreased by −3 dB (black dashed line). ***B*** Two-dimensional representation of the matrix used in the *τ*_*m*_ correction procedure. Depending on the average duration of all up-states within one voltage level, the initial 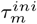 was corrected by the appropriate value to obtain the final *τ*_*m*_ estimate (see Methods). The black dashed line and the marker represent the data depicted in *A*. ***C*** Up-state vs. down-state *τ*_*m*_ for all neurons regardless of the cell type or physiological condition; the dashed line represents equality. It is clear that *τ*_*m*_ in the up-states is smaller than in the down-states, indicating a high-conductance regime due to synaptic bombardment, similar to that in neocortical neurons (Paré et al., 1998; Destexhe et al., 1999; Léger et al., 2005). ***D*** There is no significant difference in up-state *τ*_*m*_ between dMSNs and iMSNs, either in control or 6OHDA conditions. This suggests that the differences in up-state membrane power are not the result of differences in membrane dynamics between dMSNs and iMSNs. In down-states, 6OHDA dMSNs had higher *τ*_*m*_ than the control cells (*p* = 0.048). Data are shown as mean ±SEM. Control dMSNs and iMSNs are in red and blue, respectively, whereas 6OHDA dMSNs and iMSNs are in light red and light blue, respectively. \**p* < 0.05

The smoothing of the average PSD estimate introduces a shift of the −3 dB point, leading to an erroneous estimation of the effective membrane time constant. The magnitude and sign of the error depend, in a non-linear fashion, on the width of the Gaussian kernel used for smoothing in the frequency domain, and the duration of the original signal in the time domain. To account for this error, we numerically determined a correction term 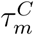, which we could then add to the initially estimated value 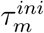, to obtain the final MSN membrane time constant estimate *τ*_*m*_. This correction term was calculated as follows. We constructed multiple surrogate “neuronal” time series by filtering Gaussian white noise signals of different durations through a set of low-pass Butterworth filters (third order, zero-phase) with predetermined cutoff frequencies. Thus, for each of the surrogate time series we knew the actual time constant (*τ*^*actual*^) of the underlying low-pass filter. We then proceeded to make an initial estimate of the time constant (*τ* ^*ini*^) as described above, using a single fixed value for the kernel width of the Gaussian smoothing function (*k*_*w*_ = 12). The error term was then defined as *τ* ^*C*^ = *τ*^*actual*^−*τ* ^*ini*^. Using this approach, we obtained the correction term *τ* ^*C*^ for signals of different durations and filters with different time constants. Next, we defined *τ* ^*C*^ as a function of *τ* ^*ini*^ and signal duration (Fig. 3*B*) to obtain the correction term 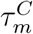 for our estimates of the MSN membrane time constant. Finally, the effective MSN membrane time constant *τ*_*m*_ was determined by adding the corresponding 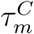 to the initial estimate 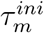.

The main weakness of this method stems from the necessity of averaging the spectral data over many trials of sufficient duration and power. That is, for the most precise estimation, the trials (states) should preferably be at least 250 ms long, and the input to the neurons should have rich enough frequency content to uncover the membrane cutoff frequency (comparable to injecting white noise into the *in vitro* recorded neuron).

Due to the underlying approximations and limitations of the method, the estimated values of the membrane time constants should not be treated as actual, precise values of those neurons’ *τ*_*m*_. Nevertheless, our approach does return consistent and comparable results across different cells when applied to the recorded data. Moreover, our analysis employing the *τ*_*m*_ estimation procedure uncovers differences in membrane time constant of the down-states similar to those previously reported in Ketzef et al. (2017) (Fig. 3*D*).

#### Spike-triggered average (STA) calculations

For every recorded neuron that spiked we extracted 12 ms of the pre-spike voltage traces. The duration of this particular time window was chosen as it roughly represents the average membrane integration window for synaptic input in the up-states, based on the estimations of the effective membrane time constants across different cell groups (Fig. 3*D*). For every cell, spikes were identified in the voltage trace, and the intervals from 0.25 ms before to 5 ms after the spike events were removed from the trace. The spiking threshold was then determined as the largest fluctuation of the first derivative of the remaining trace. The times when the derivative of the full trace crossed the threshold were taken as spike onset times. For the purpose of calculating the average of these pre-spike voltage traces (STA), we did not include any spikes occurring during state transitions, that initiated less than 12 ms after the start of an up-state, or which were occurred earlier than 17 ms after the previous spike in the same up-state. The remaining pool of spikes was divided into those that were the result of spontaneous neuronal activity and those that arose as the consequence of whisker stimulation. If, after these selections. the pool contained at least three spikes, the STA was calculated.

The STAs were compared using a permutation test. For each group comparison we collected all the cell-average traces into a single pool, shuffled their indices, and generated randomized groups by drawing as many traces from this common shuffled pool as the original groups had. For each such generated randomized group, we constructed a grand-average STA. We repeated this process 1,000 times. Significance lines were determined as the 2.5 % and 97.5 % of the voltage distributions of the random grand-average traces for each time point. The range of voltage distributions differed between groups when the number of traces belonging to randomized groups for a single comparison was different (e.g., for the comparison of spontaneous vs. evoked iMSN STAs, we had 11 spontaneous and 4 evoked cell-average traces). This difference is reflected in the voltage ranges depicted in the graphs, but it does not affect the validity of the permutation test.

#### Statistical methods

nless noted otherwise, the data are presented as mean ±SEM and were tested for normality using the Shapiro-Wilk test. Normally distributed data were tested by the unpaired two-sample Student’s t-test, and non-normally distributed data by the Wilcoxon rank-sum test (ttest2 and ranksum in MATLAB, respectively). The significance level *α* was set to 0.05. In the case of PSD comparison over different frequency bands (Fig. 2*D*), the results were corrected for multiple testing by the Holm-Bonferroni correction (Holm, 1979), and both the corrected *α*-level (*α*_*HB*_) and the calculated p-value are reported.

All data analyses were performed using custom scripts written in MATLAB R2016a (Mathworks, Inc.).

## Results

To estimate the relative strength of excitatory synaptic inputs to striatal neurons, we obtained *in vivo* whole-cell patch clamp recordings of MSNs from the dorsolateral striatum in control (dMSN *n* = 26, iMSN *n* = 18, total *n* = 44) and 6OHDA lesioned mice (dMSN *n* = 14, iMSN *n* = 9, total *n* = 23). We used optogenetic stimulation to classify MSNs online during the recording session as belonging to either the direct or indirect pathway using the optopatcher (Katz et al., 2013). Both dMSNs and iMSNs showed slow-wave membrane potential oscillations (up- and down-states), characteristic of neurons recorded in animals under ketamine-induced anesthesia (Wilson and Kawaguchi, 1996) (Fig. 2*A*). During the up-state, MSNs receive barrages of excitatory inputs from the neocortex and the thalamus. We therefore analyzed the variance and spectrum of the up-state membrane potential traces to assess the respective synaptic inputs to the two MSN types.

### dMSNs have higher spectral power in up-state than iMSNs

If a neuron soma is treated as a simple linear integrator, the mean and variance of the subthreshold membrane potential fluctuations is primarily determined by the firing rate, the number of excitatory and inhibitory inputs to a given cell, and their synaptic strength (Kuhn et al., 2004):

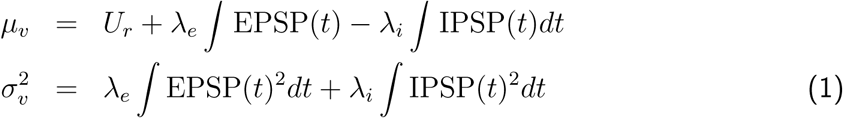

where *µ*_*v*_ and 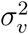 are the voltage-dependent mean and variance of the membrane potential, *U*_*r*_ is the resting membrane potential, *λ*_*e*_ and *λ*_*i*_ are the rates of excitatory and inhibitory inputs, respectively, and *EPSP*(*t*) and *IPSP*(*t*) describe the temporal shape of excitatory and inhibitory post-synaptic potentials.

From Eq. 1 it is clear that excitatory and inhibitory inputs have an opposite effect on the mean of the membrane potential of a cell receiving synaptic input. By contrast, because the calculation of the variance involves the square of the PSPs kernel, an increase in either excitatory or inhibitory inputs always results in an increase of the variance of the membrane potential (Eq. 1). Against this background, consider two neurons, *n*_*s*_ and *n*_*w*_, receiving inputs via stronger and weaker synapses, respectively. The excitatory and inhibitory inputs to these two neurons can be tuned such that both *n*_*s*_ and *n*_*w*_ have the same mean membrane potential. However, due to the stronger synaptic weights and, hence, larger post-synaptic potentials, the neuron *n*_*s*_ will exhibit a larger membrane potential variance than the neuron *n*_*w*_. This example illustrates that the mean membrane potential is not an adequate measure for the overall synaptic input, but by comparing the variances it is possible to determine if two neurons receive different amounts of synaptic inputs. This requires that the two neurons receive uncorrelated synaptic inputs and that their membrane time constants are similar.

Since the variance in time-domain equals the power spectral density (PSD) in frequency domain (Parseval’s theorem), the PSD gives an estimate of the variance for every frequency in the signal (Papoulis and Pillai, 2002). Therefore, we measured the PSD of the membrane potential for every detected up- and down-state of a cell (Fig. 2*A*, see Methods).

For each MSN type we constructed a grand-average PSD estimate for both control and 6OHDA conditions (Fig. 2B). Direct comparison of these grand-averages revealed that dMSNs had consistently higher PSD than iMSNs over all examined frequency bands under control conditions. In particular, in three prominent, higher-frequency bands (*β*: 13-30 Hz, *Z* = 2.47, *p* = 0.0135, *α*_*HB*_ = 0.0167; low-*γ*: 30-70 Hz, *t*_(42)_ = 2.72, *p* = 0.0095, *α*_*HB*_ = 0.01; and high-*γ*: 70-150 Hz, *Z* = 2.57, *p* = 0.01, *α*_*HB*_ = 0.0125) dMSNs showed significantly higher power than iMSNs (Fig. 2*C*, top left). Because the total power spectral density of the membrane potential in a selected frequency band equals the variance of the membrane potential in that frequency band (Papoulis and Pillai, 2002), the heightened power of dMSN up-state membrane potentials in control animals is indicative of stronger voltage fluctuations as compared to iMSNs. Unlike under control conditions, in the DA-depleted striatum we found no difference in the spectral power of up-state membrane potential fluctuations of dMSNs and iMSNs (across all bands *p* > 0.68, Fig. 2*C*, top right). Comparison of the up-state control vs. DA-depleted conditions revealed a significant difference in dMSN *α*-band (8-13 Hz) power (*Z* = 2.62, *p* = 0.0087, *α*_*HB*_ = 0.01). DA depletion did not affect the spectral power of iMSNs (control vs. DA-depleted conditions across all bands *p* > 0.55). Finally, in down-states, there was no significant difference in spectral power of dMSNs and iMSNs in either condition (the calculated significance value was always above the corrected alpha level).

Given that up-states are thought to be primarily synaptically driven (Wilson and Kawaguchi, 1996; Stern et al., 1997), our results indicate that the increased power of dMSNs, especially in the higher-frequency bands, compared to iMSNs in the control case stems from stronger total input to direct pathway striatal neurons. Furthermore, our results suggest that in dopamine-depleted conditions, the total input to dMSNs is either significantly reduced and/or is more similar to the input to iMSNs.

### MSN membrane time constant does not underlie the differences in high-frequency power

The difference in high-frequency power between dMSNs and iMSNs may be caused by a difference in the time constants of the two neuron types. We estimated the effective time constant using the spectrum of the membrane potential fluctuations (see Methods).

We found that the effective time constants for both dMSNs and iMSNs in the up-states were smaller than in the down-states (Table 1, Fig. 3*C*). On average, the ratio of down-state to up-state effective membrane time constant across all groups was 1.86 (Fig. 3*D*, dMSN control 1.76, iMSN control 1.85, dMSN 6OHDA 2.05, iMSN 6OHDA 1.77). This is similar to the case of neocortical neurons, which also show a shorter time constant in up-states (Paré et al., 1998; Destexhe et al., 1999; Léger et al., 2005). However, in MSNs this ratio is not as large as has been reported for neocortical neurons (Reig and Silberberg, 2014), presumably because of the closing of potassium inward rectifier (Kir) channels in MSNs, happening as the membrane depolarizes (Waters and Helmchen, 2006; Nisenbaum and Wilson, 1995).

**Table 1.**
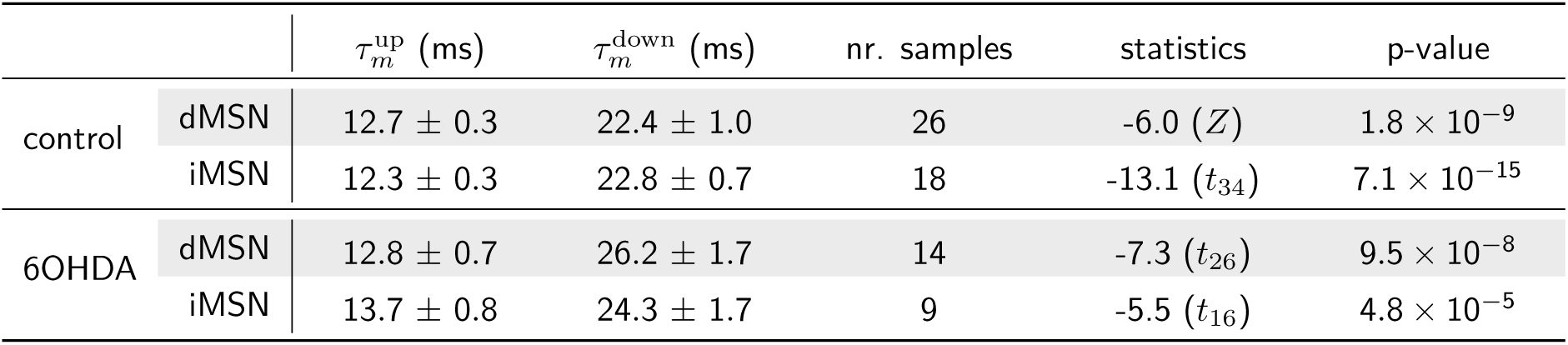
Comparison of the effective time constants of dMSNs and iMSns in the up-states vs. down-states. For both MSN types, across both healthy and dopamine-depleated conditions, up-states exhibited significantly faster membrane dynamics than down-states.

Further comparisons showed no significant difference between the up-state effective time constants of dMSNs and iMSNs in control or 6OHDA conditions (in all cases *p* > 0.085; Fig. 3*D*). However, in the down-states, the effective *τ*_*m*_ of dMSNs was slightly larger in the 6OHDA condition than in the control (*t*_(38)_ = −2.05, *p* = 0.048; control *n* = 26, 6OHDA *n* = 14), whereas no such difference was present for the iMSNs (*p* = 0.37; Fig. 3*D*). These results are partially consistent with previously reported measurements of input resistance using standard methods in MSN down-states (Ketzef et al., 2017).

Taken together, these results clearly suggest that the differences in the power spectra of up-state sub-threshold membrane potential fluctuations between dMSNs and iMSNs (Fig. 2*D*) are not the result of different membrane time constants of the two types of neurons. Moreover, the lower membrane time constant of MSNs in the up-state suggests that these neurons also operate in a relatively high conductance regime.

### dMSNs receive stronger input from mouse sensory cortex than iMSNs

In the analysis so far we focused on subthreshold membrane potential sections, in which neurons did not fire action potentials during the up-states. To further test the hypothesis that dMSNs indeed receive stronger inputs than iMSNs, we investigated the membrane fluctuations leading to action potential discharges. To this end, we obtained the spike-triggered average (STA) of the membrane potential immediately preceding action potential discharge for each neuron (Fig. 3*A-C*). If dMSNs would indeed receive stronger inputs, we would expect the corresponding STA traces to approach the spike threshold with a steeper slope, compared to the STA traces of iMSNs. To better quantify this difference, we sub-divided the spikes of each neuron into those corresponding to spontaneous spiking activity and those evoked by whisker deflections with brief air puffs. This STA analysis was only performed for healthy animals, as in our data MSNs recorded from DA-depleted mice elicited only a very small number of spikes, not sufficient for STA analysis.

Comparison of the grand-average STAs for the two MSN types upon bilateral whisker stimulation revealed that dMSNs indeed depolarized to the spike threshold much faster than iMSNs. For the spontaneously generated spikes, this applied as well, although the difference was less prominent. Comparing the average membrane potential 12 ms before spike time (the duration of the average up-state integration window shown in Fig. 3*D*), we found that dMSN membrane potentials were on average 1.3 mV more hyperpolarized than those of iMSNs, resulting in steeper depolarization slopes (*k*) preceding spike initiation (Fig. 4*D*, *t* = −12 ms; dMSN, −6.27±0.34 mV, *k* = 0.31 mV/ms, *n* = 10; iMSN, −4.97±0.53 mV, *k* = 0.28 mV/ms, *n* = 10). This difference was even bigger for whisker stimulation evoked spikes (Fig. 4*E*, *t* = −12 ms; dMSN, −9.95±1.16 mV, *k* = 0.58 mV/ms, *n* = 5; iMSN, −6.01±0.72 mV, *k* = 0.34 mV/ms, *n* = 4).

**Fig 4.**
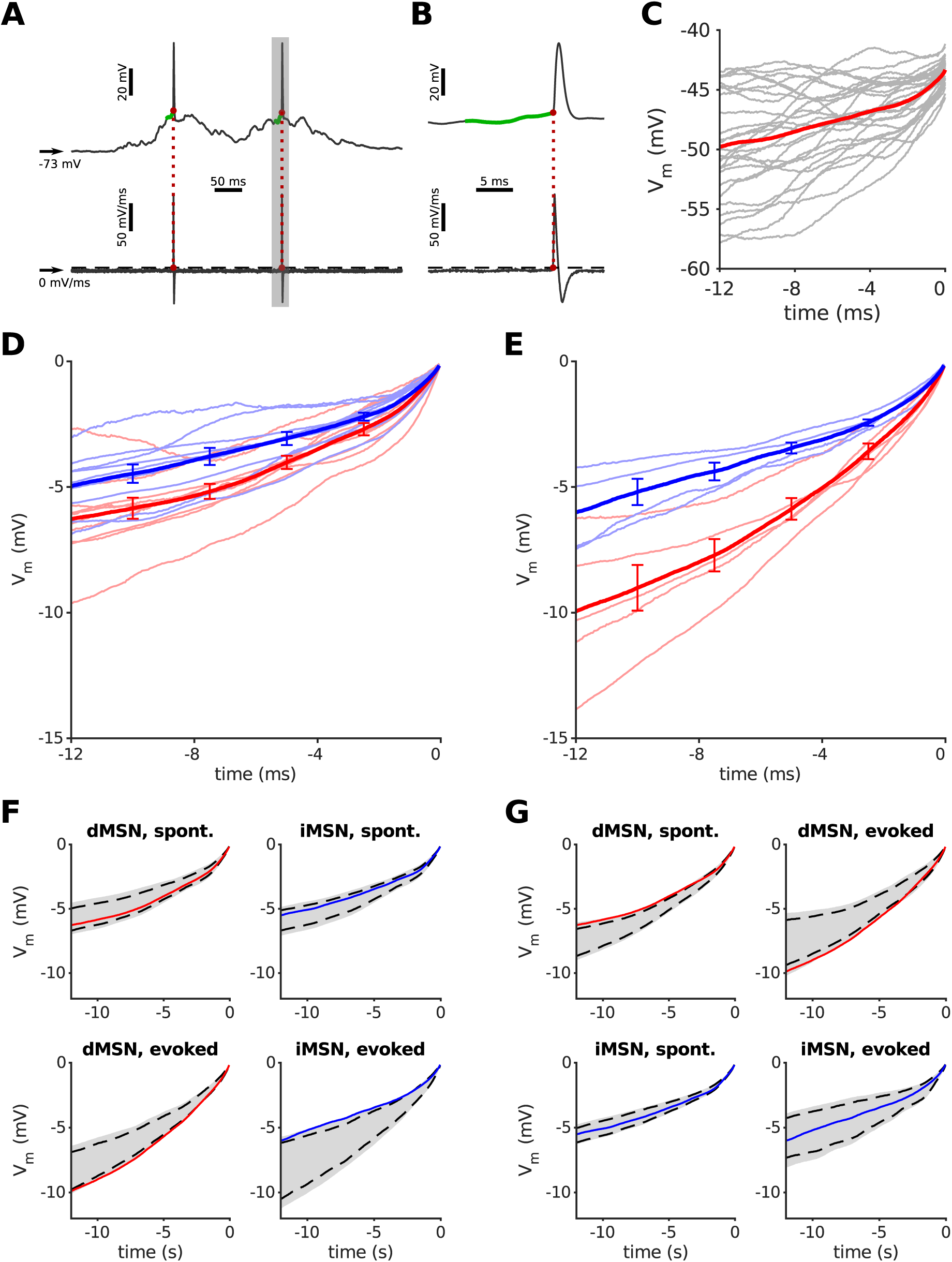
dMSNs accelerate faster towards firing threshold than iMSNs when receiving input from barrel cortex. ***A*** Example of the estimation of the firing threshold and extraction of the pre-spike voltage trace. *Top*: membrane potential of a dMSN in the up-state; *bottom*: its first-order derivative *dV /dt*. The spiking threshold was determined as the highest voltage deflection seen in the derivative that didn’t produce a spike (black dashed line); when the derivative crossed the threshold, this marked the start of an action potential (red dot). The voltage trace during 12 ms preceding the spike onset is marked in green. ***B*** Expanded view of the shaded area in *A*. ***C*** Example of calculating the spike-triggered average (STA, red curve) for the neuron in *A*. Gray traces are 12 ms pre-spike intervals from individual up-states producing a spike. ***D*** Comparison of grand-average STAs of dMSNs and iMSNs in control conditions (thick lines in red and blue, respectively) when action potentials were generated by spontaneous activity. Faint red and blue traces show STAs for individual neurons of corresponding MSN types. All traces were aligned to spike onset. Error bars represent SEM. ***E*** Same as in *D*, but the action potentials were generated by whisker stimulation and synaptic input from the barrel cortex. Note that the grand-average STA of dMSNs is accelerating faster toward spike onset, indicating stronger synaptic input to these neurons. ***F*** The permutation test shows that evoked dMSN and iMSN STAs differed significantly, by falling in the bottom and top 2.5 % of voltage distributions, respectively. However, no such difference was observed for spontaneous traces. ***G*** While there was no marked difference between spontaneous and evoked iMSN STAs, dMSN traces differed significantly across the two conditions.

We further examined the STA differences between dMSNs and iMSNs by utilizing a permutation test (see Methods). When a grand-average STA trace would fall above 97.5 % or below 2.5 % voltage distribution line, we deemed that result significant. We found that the comparison of spontaneous dMSN and iMSN STAs yielded no significant difference. However, the evoked STA traces between dMSNs and iMSNs were markedly different (Fig. 4*F*). Furthermore, the additional input from the sensory cortex seems to have specifically targeted dMSNs, as their STA traces varied significantly between the spontaneous and evoked conditions, whereas no major change was observable for the same comparison of iMSNs (Fig. 4*G*).

Thus, the results of our STA analysis also support the notion that dMSNs receive stronger synaptic input than iMSNs in healthy animals.

## Discussion

Here we provided evidence that *in vivo* dMSNs receive stronger synaptic input than iMSNs and that this difference is attenuated in dopamine-depleted animals. These findings were based on two observations: (1) dMSNs showed significantly higher spectral power than iMSNs in the up-states, especially in the higher-frequency bands (Fig. 2*C,D*), and (2) in both spontaneous and stimulus-induced spikes dMSNs membranes depolarized faster than iMSNs before reaching spike-threshold, as revealed by their STAs (Fig. 4). These results provide support for the theoretical prediction that direct-pathway MSNs in healthy state animals receive stronger synaptic input than iMSNs (Bahuguna et al., 2015). In addition, we showed by spectral analysis that the effective membrane time constant of MSNs during up-states is significantly shorter than in down-states, indicating that synaptic inputs affect the membrane conductance to a larger extent than hyperpolarization-activated conductances mediated by the Kir channels in MSNs.

Paired recordings in slices revealed that iMSNs form more and stronger synaptic connections onto dMSNs than they receive from them (Taverna et al., 2008; Planert et al., 2010). In addition, fast spiking interneurons also form more connections onto dMSNs than onto iMSNs (Planert et al., 2010; Gittis et al., 2010). Given these differences in connectivity, Bahuguna et al. (2015) predicted that dMSNs must receive more or stronger excitatory inputs if both dMSNs and iMSNs were to be co-activated (Cui et al., 2013) or have comparable activity levels in both ongoing and stimulus evoked activity (Sippy et al., 2015), as has been observed experimentally. Consistent with this prediction, Parker et al. (2016) showed that, in healthy animals *in vitro*, dMSNs receive stronger excitatory input from both thalamo-striatal and cortico-striatal projections. They also showed that in dopamine-depleted mice thalamo-striatal inputs to iMSNs become stronger than to dMSNs, whereas cortico-striatal projections remain largely unchanged.

Here, we show that the disparity between the excitatory inputs to dMSNs and iMSNs is also maintained *in vivo* in the up-states, which closely resemble the awake state of an animal (Destexhe et al., 2003; Haider et al., 2012). In our analysis we assumed that larger fluctuations in the membrane potential are a reflection of stronger synaptic weights and/or correlated inputs. Because both thalamus and cortex are co-activated in the up-states, we cannot distinguish between thalamo-striatal and cortico-striatal inputs. However, by selectively silencing thalamic inputs to the striatum (using optogenetic or chemogenetic approaches) it should be possible to determine the relative contributions of thalamo-striatal and cortico-striatal inputs *in vivo* following our approach. Another major limitation of our analysis is that we cannot separate excitatory from inhibitory inputs. In fact, an increase in either type of synaptic inputs can increase the membrane potential variance (Kuhn et al., 2004). However, our comparison of STAs suggests that dMSNs are more likely to receive stronger excitatory inputs because during both, spontaneous and stimulus-evoked activity, dMSNs depolarize faster to the action-potential threshold than iMSNs (Fig. 4).

The size of membrane potential fluctuations is affected by the mean membrane potential which determines the synaptic driving force and the membrane time constant (Kuhn et al., 2004). In our data we did not find a significant difference between the mean up-state membrane potentials of dMSNs and iMSNs in either control or 6OHDA conditions (in all cases *p* > 0.138, data not shown), and no significant difference in up-state *τ*_*m*_ estimations either (Fig. 3*D*). The latter is contrary to previous findings *in vitro*, where measurements of whole-cell capacitances and input resistances of dMSNs and iMSNs suggest that their membrane time constants are different (Gertler et al., 2008; Fieblinger et al., 2014). It should be born in mind, however, that under *in vivo* conditions the membrane properties are affected by the ongoing synaptic activity (Kuhn et al., 2004) and that synaptic inputs can easily overcome the differences in the neuron membrane properties measured *in vitro* (Destexhe et al., 2003, 2007).

Furthermore, our data shows that in MSNs the effective membrane time constant in the up-states is on average 46 % smaller than in the down-states (Table 1, Fig. 3*C*), indicating a high-conductance state in striatal neurons in the presence of synaptic inputs, similar to that of neocortical neurons (Destexhe et al., 2003; Léger et al., 2005; Destexhe et al., 2007). That is, in the up-states, the membrane time constant is strongly influenced by synaptic inputs.

Finally, our results also provide new insights into how dopamine affects the excitatory inputs to the MSNs. We found that in dopamine-depleted animals both dMSNs and iMSNs receive similar amounts of excitatory inputs. These results are consistent with our previous findings that in healthy animals dMSNs exhibit stronger response to contralateral sensory stimulation than iMSNs, and that these differences are diminished in dopamine-depleted mice (Reig and Silberberg, 2014; Ketzef et al., 2017). Thus, our results and previous findings (Parker et al., 2016; Ketzef et al., 2017) suggest that dopamine is important to maintain the difference in the excitatory inputs to dMSNs and iMSNs. In the absence of dopamine, excitatory inputs to dMSNs are weaker and the indirect pathway is active for a much larger range of cortico-thalamic inputs, resulting in dysfunctional action selection and action initiation (Bahuguna et al., 2015). While we have provided evidence for stronger total synaptic input to the dMSNs as compared to iMSNs, it is still unclear whether the extra excitation to the dMSNs is due to stronger cortical and/or thalamic inputs. Furthermore, it also not clear whether the larger membrane potential fluctuations in dMSNs are due to stronger synapses or to higher input correlations. More dedicated experiments involving selective correlated activation of cortical and thalamic neurons will help resolving these questions.

## Acknowledgements

We thank Dr. Jyotika Bahuguna, Dr. Rita Almeida, Matthijs Dorst, Michael Zohar and Yvonne Johansson for their technical support and advice. This work was funded in parts by: the EU Erasmus Mundus Joint Doctorate Program ‘EUROSPIN’, The International Graduate Academy (IGA) of the Freiburg Research Services (to Marko Filipović), Swedish Research Council (Research Project Grant, StratNeuro), Parkinsonfonden grant (to Arvind Kumar), Swedish Research Council (Research Project Grant) (to Gilad Silberberg), the German Research Foundation (DFG#1086 BrainLinks BrainTools) (to Ad Aertsen and Arvind Kumar).

